# OBESITY-INDUCED ENDOTHELIAL FENESTRATION AND CAPILLARY LEAKAGE CONTRIBUTE TO INCREASED PAIN SENSATION

**DOI:** 10.64898/2026.03.13.711502

**Authors:** Yuta Koui, Jonathan Richard Shin, Shuxuan Song, Christian A. Combs, Yoh-suke Mukouyama

## Abstract

Peripheral pain sensation is regulated by interactions between sensory nerves and various tissue cells. In obese patients with painful small fiber neuropathy, skin sensory nerves are often hypersensitive. While obesity is known to cause circulation-related vascular abnormalities, how these changes affect sensory dysfunction is not fully understood. In this study, we found that in a diet-induced obesity mouse model, skin capillaries become fenestrated, allowing insulin to diffuse into the avascular epidermis. This exposure triggers the production and secretion of nerve growth factor (NGF) from epidermal keratinocytes via insulin signaling with the forkhead box O1 (FOXO1) transcription factor. Elevated NGF leads to heightened sensory hypersensitivity by enhancing transient receptor potential vanilloid subtype 1 (TRPV1) in sensory nerves. Controlling capillary permeability reduces abnormal NGF expression and attenuates pain hypersensitivity. These findings nominate peripheral nerve-associated capillary permeability as a novel therapeutic target in obesity-associated sensory dysfunction.

## Introduction

Obesity can cause neuropathic pain, characterized by hypersensitivity in sensory axons in the skin, a condition known as painful small fiber neuropathy, while patients with severe diabetes often experience a loss of sensitivity due to diabetic neuropathy ^1,2^. Additionally, systemic hyperglycemia, dyslipidemia, insulin resistance, and/or elevated blood pressure associated with obesity contribute to cardiovascular diseases, including vascular abnormalities in various tissues ^3,4^. However, the impact of these vascular abnormalities on skin sensory dysfunction remains poorly understood.

Our previous studies with diet-induced obesity (DIO) mice established an association between obesity and enhanced pain behavior including sensory hypersensitivity in skin sensory nerves ^5^. We found that DIO mice fed a high-fat diet for 16 weeks, which were starting at 6 weeks-of-age, exhibited enhanced pain behaviors when exposed to the irritant capsaicin at 22 weeks-of-age ^5^. Epidermal keratinocytes in the skin of DIO mice produced and secreted nerve growth factor (NGF) in response to elevated insulin levels, which sensitized sensory axons through NGF-Tropomyosin receptor kinase A (TrkA)-phosphatidylinositol 3-kinase (PI3K) signaling pathway ^5^. In contrast, DIO mice at 30 weeks-of-age (with an additional 8 weeks of high-fat diet) showed relatively reduced pain behaviors and skin sensory activity, suggesting that a temporal window for painful small fiber neuropathy is followed by painless neuropathy during the progression of pre-diabetic obesity. Interestingly, we found elevated levels of the vascular permeability marker, endothelial cell (EC)-specific protein plasmalemma vesicle-associated protein (PLVAP) ^6–9^ in dermal vessels that correlated with the temporal window of neuropathic pain in DIO mice at 22 weeks-of-age ^5^. These findings led us to hypothesize that increased vascular permeability contributes to sensory dysfunction in DIO mice.

To address this, we sought to directly manipulate PLVAP-mediated vascular barrier function in DIO mice, and characterize associated pain behaviors. We reported severe abnormalities in skin vascular barrier integrity along with increased PLVAP expression and capillary fenestration. Application of a neutralizing anti-PLVAP antibody reduced vascular permeability, leading to decreased pain behavior and sensory hypersensitivity. Taken together with our observation that insulin signaling was diminished and NGF expression was reduced in the epidermal keratinocytes of the antibody-treated DIO mice, our findings suggest that inhibiting vascular permeability limits insulin diffusion from the skin capillaries into the epidermis. This reduced insulin availability ultimately suppresses insulin-mediated NGF production in the epidermal keratinocytes and NGF-TrkA signaling in keratinocyte-associated sensory axons. Overall, our findings highlight how increased vascular permeability contributes to sensory dysfunction in painful small fiber neuropathy associated with obesity, and opens important new treatment targets for this disorder.

## Results

### Superficial skin capillaries become fenestrated in DIO mice

To characterize vascular architectural changes in DIO mice, we performed whole-mount immunostaining of ear skin using the pan-EC marker platelet endothelial cell adhesion molecule 1 (PECAM-1). Consistent with previous studies showing arteries and veins in the deep dermis, branched arterioles and venules in the intermediate dermis, and capillaries in the superficial dermis ^10^, we documented here a hierarchical vascular branching network spanning the entire skin dermis, but excluding the epidermis (Figures 1A-1C). Since superficial skin capillaries limit molecular exchange between blood and tissue by forming a barrier endothelium ^11,12^, they typically do not form fenestrae, which are window-like open pores ^7^. To investigate structural changes in superficial dermal capillaries due to obesity, we utilized a diet-induced obesity (DIO) mouse model (Figure 1D) fed a high-fat diet (60% fat, 20% protein, and 20% carbohydrate kcal) for 16 weeks, starting at 6 weeks-of-age. At 22 weeks-of-age, we observed significant increase in the body weight, blood glucose levels, and serum insulin levels in DIO mice, compared to control mice fed a normal diet (10% fat, 20% protein, and 70% carbohydrate kcal) ^5^. Our previous studies in these mice revealed enhanced pain behaviors and sensory hypersensitivity when exposed to capsaicin ^5^. Whole-mount immunostaining of ear skin using PECAM-1 and PLVAP (clone MECA-32), along with the vascular smooth muscle cell marker alpha smooth muscle actin αSMA, demonstrated increased PLVAP expression in PECAM-1^+^/αSMA^−^ capillaries located in the superficial dermis of DIO mice, compared to control mice (Figures 1E and 1F). It is noteworthy that PLVAP expression is restricted to ECs, indicating its specificity to these cells. Transmission electron microscopy confirmed that over 20% of superficial dermal capillary ECs in DIO mice formed fenestrae, whereas control capillary ECs did not (Figures 1G and 1H). These findings suggest that the superficial dermal capillaries in DIO mice exhibit leaky barrier structures due to abnormal formation of endothelial fenestrations. Vascular permeability is regulated not only by the formation of fenestrae in ECs but also by the stability of EC-pericyte association ^13^. We confirmed that there were no significant changes in adhesion junctions between ECs (Figures S1A and S1B) and platelet-derived growth factor receptor beta^+^ (PDGFRß^+^) pericyte coverage associated with PECAM-1^+^ capillaries (Figures S1C and S1D). These findings indicate that DIO promotes formation of endothelial fenestration in the superficial dermal capillaries without affecting the stability of EC-adhesion junctions and pericyte coverage.

**Figure 1.**
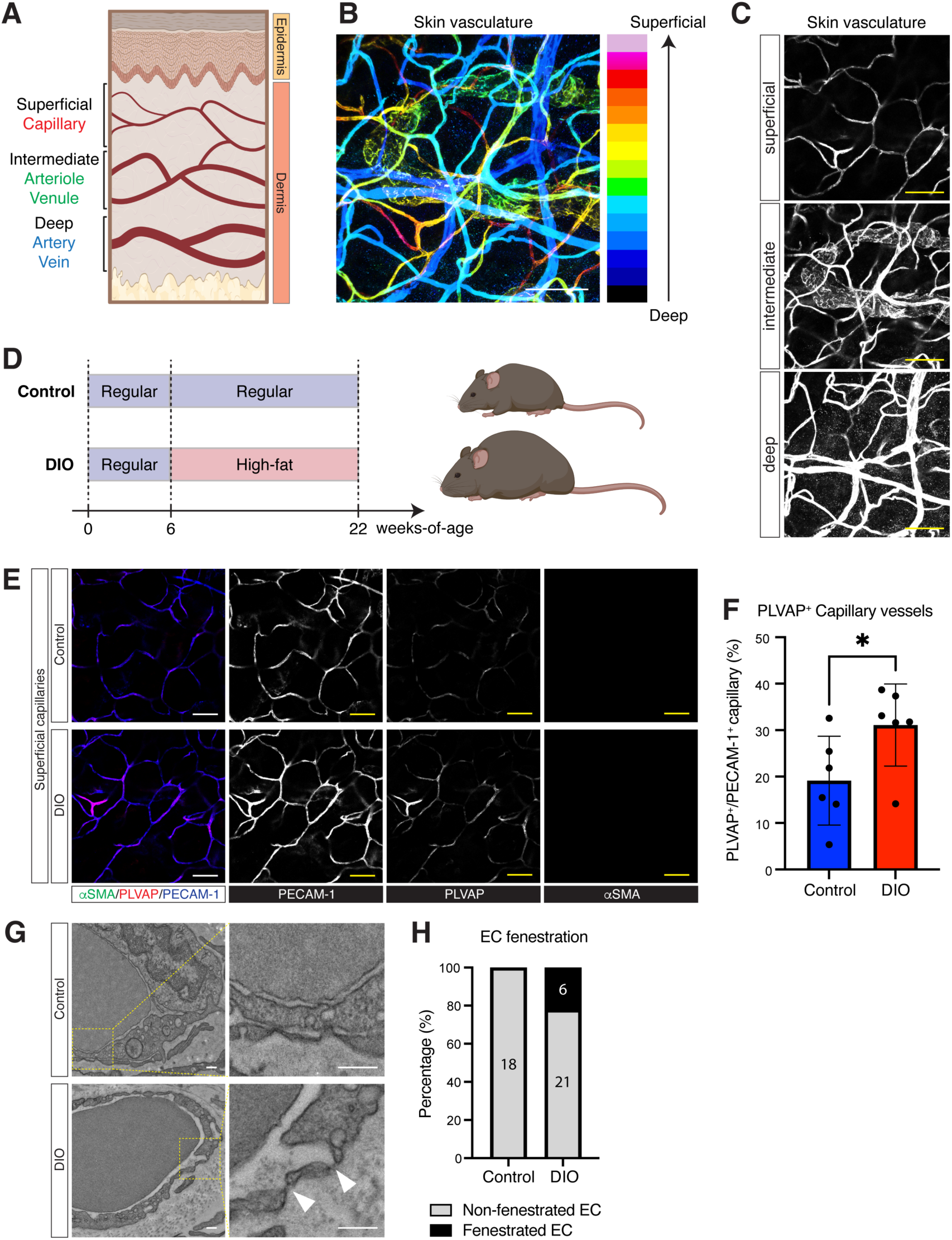
Superficial skin capillaries become fenestrated in DIO mice. (A) Schematic diagram of skin vasculature illustrates the organization of blood vessels in the skin layers. In the deep dermis, there are large-diameter blood vessels, including arteries and veins. The intermediate dermis contains arterioles and venules that branch from these arteries and veins, forming an intricate network. Lymphatic vessels are also located in the intermediate dermis. In the superficial dermis, capillaries form a highly branched network. (B) A representative maximum projection image from a whole-mount immunohistochemical analysis of mouse ear skin is presented. This analysis uses the pan-endothelial cell (EC) marker PECAM-1 to visualize skin vasculature from the deep to the superficial dermis. Different colors in the image indicate varying depths (Z-depth) within the dermis. Scale bar: 100 μm. (C) Representative whole-mount images of PECAM-1^+^ vasculature in the superficial, intermediate, and deep dermis of mouse ear skin are shown. Scale bars: 100 μm. (D) Experimental outline for generating diet-induced obesity (DIO) mice. Mice were fed either regular diet (10 Kcal % fat) or high-fat diet (60 Kcal % fat) from 6 weeks-of-age to 22 weeks-of-age. (E) Representative whole-mount images of the superficial dermal vasculature in the ear skin of control and DIO mice at 22 weeks-of-age, labeled with antibodies for the vascular smooth muscle cell marker αSMA (green or white), the vascular permeability marker PLVAP (MECA-32, red or white), along with PECAM-1 (blue or white) are presented. Scale bars: 100 μm. (F) Quantification of PLVAP^+^/PECAM-1^+^ capillaries in the superficial dermal vasculature from control and DIO mice is shown. The sample size is N = 6 in each group. (G) Representative transmission electron microscopy images of capillary ECs in the superficial dermis from control and DIO mice are presented. The dotted box regions in the left panels are magnified in the right panels. Fenestrae were observed only in DIO capillary ECs (arrowheads). Scale bars: 200 nm. (H) Quantification of endothelial fenestration in control and DIO capillary ECs is provided, showing both the number and percentage of non-fenestrated and fenestrated ECs. The sample sizes are as follows: N = 18 in control, N = 27 in DIO. Results are shown as the mean ± SEM. *p<0.05. P values were determined by the parametric two-tailed t test. The schematic diagrams and graphic summary were partially created with BioRender.com.

### Administration of anti-PLVAP antibody inhibits vascular hyperpermeability in the DIO skin

PLVAP is specifically expressed in ECs as an important molecular component of fenestrae and caveolae ^6–9^, and accentuates vascular leak in laser-induced choroidal neovascularization ^14^. To establish a mechanistic role for PLVAP in DIO-induced skin vascular barrier dysfunction (Figure 2A), we continuously delivered a neutralizing anti-PLVAP antibody via subcutaneous osmotic pump to DIO mice from 20 to 22 weeks-of-age, and assessed vascular leak with a fluorescent tracer (Figure 2B). This regimen did not affect the health, body weight, blood glucose levels, or the branching structures of skin vasculature in DIO mice (data not shown). To assess vascular leakage, we used a 40 kDa fluorescent-conjugated dextran, because endothelial fenestrae regulate the leakage of molecules smaller than 50 kDa ^8^. We quantitatively measured dextran leakage in the superficial dermis over 10 minutes using intravital live imaging of ear skin (Figure 2C). Our findings revealed that compared to the ear skin of control mice, 40 kDa dextran extravasation occurred earlier in the ear skin of DIO mice. These findings indicate that formation of fenestrae in DIO capillaries enhances vascular leakage (Figures 2D-2F). Importantly, the administration of the anti-PLVAP antibody led to a significant reduction in dextran 40 kDa leakage compared to control saline administration in DIO mice, indicating that the anti-PLVAP antibody effectively inhibits vascular leakage (Figures 2D-2F).

**Figure 2.**
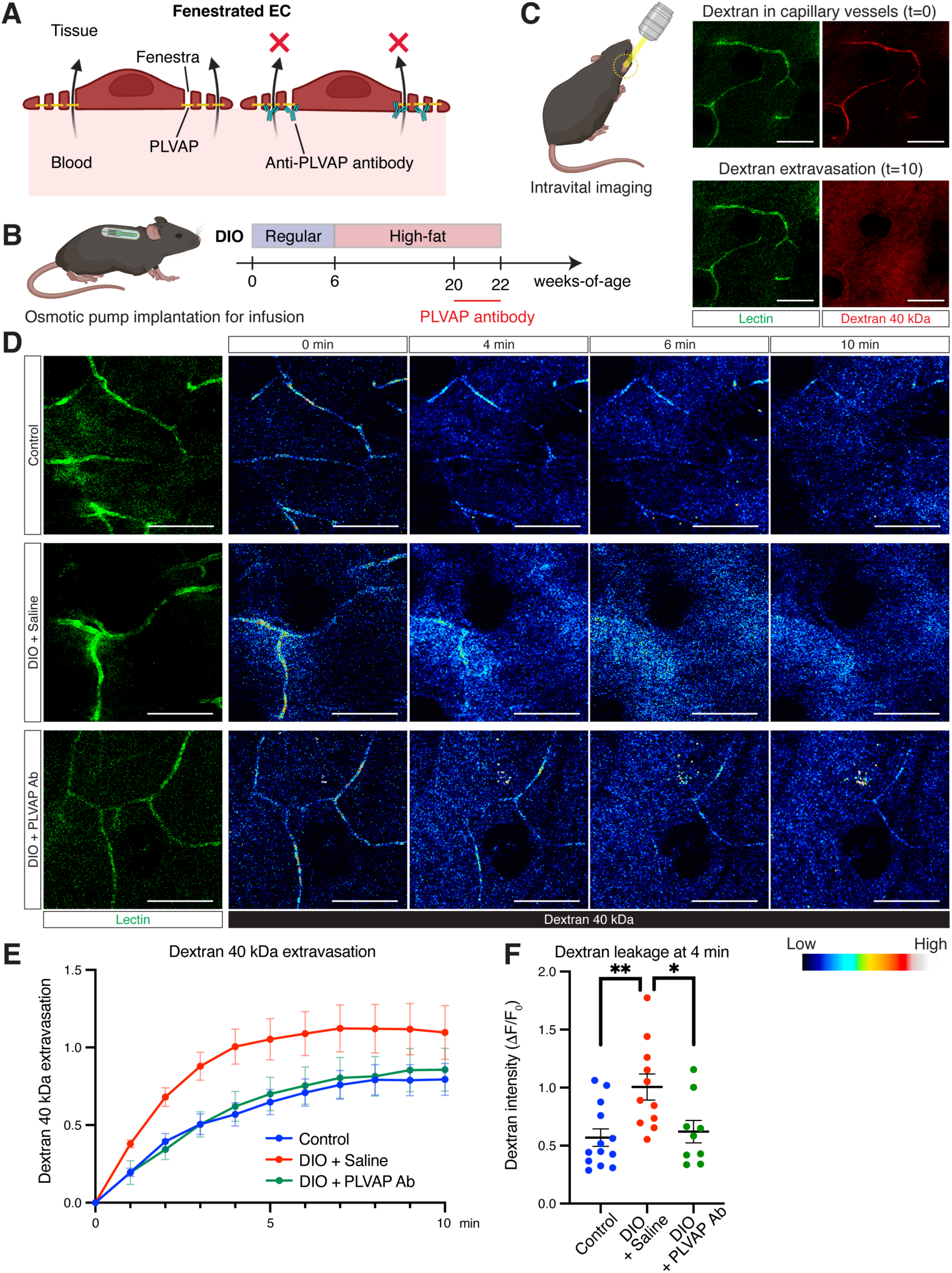
PLVAP inhibitor manages vascular hyperpermeability in DIO skin. (A) Schematic diagram illustrates capillary ECs in the superficial dermal vasculature with or without a neutralizing anti-PLVAP antibody. In DIO capillary ECs, fenestrae form, which facilitates molecular leakage from blood to tissue (left). The anti-PLVAP antibody binds to the PLVAP protein, possibly obstructing the fenestrae and inhibiting molecular leakage (right). (B) Schematic diagram illustrating the administration of the anti-PLVAP antibody into DIO mice from 20 weeks-of-age to 22 weeks-of-age using an osmotic pump. (C) Illustration shows intravital imaging of mouse ear skin, along with representative images of dextran extravasation from superficial capillaries. Lectin (green) labels capillaries, and dextran (40 kDa, red) is visible inside the capillaries immediately after injection (t=0), gradually extravasating thereafter (t=10). Scale bars: 100 μm. (D) Representative time-course images show the extravasation kinetics of dextran (40 kDa) in control skin, DIO skin treated with saline, and DIO skin treated with the neutralizing anti-PLVAP antibody. Lectin labels capillaries (green) in the superficial dermis. Time-course rainbow color images show the intensity of dextran. The amount of extravasated dextran is quantified based on the intensity of the dextran signal outside the lectin^+^ capillaries. Scale bars: 100 μm. (E) Changes in dextran (40 kDa) extravasation are shown for control skin (blue), DIO skin treated with saline (red), and DIO skin treated with the neutralizing anti-PLVAP antibody (green). (F) Quantitative measurements of dextran (40 kDa) extravasation at the 4-minute mark are shown. The sample sizes are as follows: N = 13 in control, N = 11 in DIO + Saline, N = 9 in DIO + PLVAP ab. Results are shown as the mean ± SEM. *p<0.05, **p<0.01. P values were determined by the parametric two-tailed t test. The schematic diagrams and graphic summary were partially created with BioRender.com.

### Vascular hyperpermeability causes pain behaviors and skin sensory hypersensitivity in *the DIO model*

Our previous studies showed that noxious pain stimuli evoke enhanced forelimb wiping responses and sensory hypersensitivity in DIO mice at 22 weeks-of-age (Figure 3A) ^5^. To examine whether inhibition of vascular hyperpermeability affects enhanced pain behaviors and sensory hypersensitivity in DIO skin, we implanted an osmotic pump filled with saline, an IgG control, or a neutralizing anti-PLVAP antibody into sensory neuron-specific calcium indicator mice, *Pirt-GCaMP3* mice (Figures 2B and 3B) ^15^. We applied capsaicin, an agonist for the transient receptor potential vanilloid subtype 1 (TRPV1) channel, to the skin behind the ear, and measured wiping responses as an indicator of pain behavior over a 10-minute period (Figure S2A) ^16,17^, while following associated changes in nerve calcium signaling (Figure 3C). Compared to the control mice fed a normal diet, DIO mice treated with saline showed a significant increase in wiping behaviors in response to capsaicin stimulation, matched by increased sensory nerve calcium transients. Remarkably, these enhanced wiping responses and calcium transients were almost entirely suppressed by administration of the anti-PLVAP antibody in DIO mice (Figures 3D and S2B; Figures 3E, 3F, and S2C). These findings establish that inhibition of vascular permeability attenuates pain behaviors and skin sensory hypersensitivity in the DIO model. We should note that administering an IgG control does not affect pain behavior or sensory responses to capsaicin in the DIO skin. Interestingly, reduced permeability and skin hypersensitivity was associated with enhanced intraepidermal nerve fiber (IENF) density as demonstrated by neuron-specific class III ß-tubulin (Tuj1) and keratin 14 (K14) immunostaining (Figures S2D and S2E), suggesting neuroprotective/nerve-sparing effects from chronic anti-PLVAP treatment.

**Figure 3.**
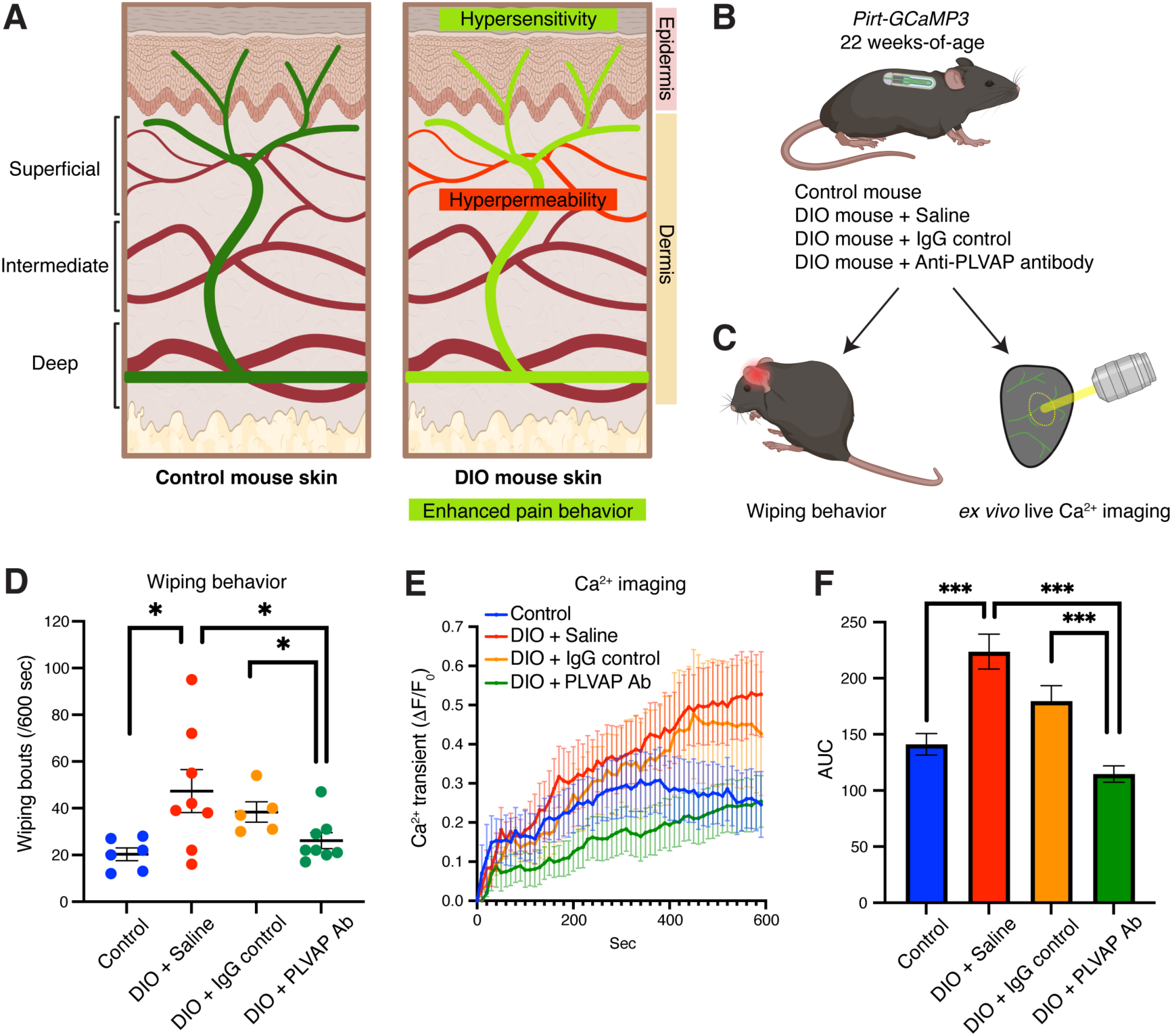
Capillary hyperpermeability leads to enhanced pain behavior and sensory hypersensitivity in DIO mice. (A) Schematic diagram illustrates the changes in vascular structure and sensory functions in the skin between control and DIO mice. In the skin of DIO mice, capillary ECs become fenestrated, leading to increased vascular permeability. Additionally, DIO mice exhibit enhanced pain behavior and sensory hypersensitivity^5^. (B) Illustration shows the implantation of an osmotic pump in sensory neuron-specific *Pirt-GCaMP3* calcium reporter mice. This pump is used to administer saline, the IgG control, or the neutralizing anti-PLVAP antibody. The sample sizes are as follows: N = 6 in *Pirt-GCaMP3* mice on a control diet (control), N = 8 in *Pirt-GCaMP3* mice with DIO receiving saline (DIO + Saline), N = 5 in *Pirt-GCaMP3* mice with DIO receiving IgG control (DIO + IgG control), N = 8 in *Pirt-GCaMP3* mice with DIO receiving the anti-PLVAP antibody (DIO + PLVAP Ab). (C) Illustrations depict the capsaicin-mediated acute pain behavior assay (left) and *ex vivo* Ca^2+^ imaging of peripheral terminals of nociceptive neurons located in the epidermis of the ear skin (right). (D) Total forelimb wiping responses following capsaicin application are shown for control, DIO + Saline, DIO + IgG control, and DIO + PLVAP Ab. (E) Quantification of Ca^2+^ responses within the ear skin of control mice, DIO mice with saline, DIO mice with the IgG, and DIO mice with the neutralizing anti-PLVAP antibody is shown. The Ca^2+^ transients were normalized by the baseline Ca^2+^ transient (ΔF/F_0_). (F) The integrated Ca^2+^ transient (ΔF/F0) was calculated as the area under the curve (AUC). Results are shown as the mean ± SEM. *p<0.05, ***p<0.001. P values were determined by the parametric two-tailed t test. The schematic diagrams and graphic summary were partially created with BioRender.com.

### Vascular hyperpermeability enhances insulin-mediated NGF production in DIO skin keratinocytes

We previously showed that keratinocyte-derived NGF promotes sensory hypersensitivity by enhancing TRPV1 receptor, likely through the NGF-TrkA-PI3K signaling axis (Figure 4A) ^5,18–23^. Indeed, short-term treatment of the DIO ear skin with a neutralizing anti-NGF antibody significantly reduced hypersensitivity ^5^. Among molecules released from leaky capillaries in DIO skin, we found that insulin exposure promotes *NGF* expression at the mRNA level in cultured human keratinocytes (Figure S3A). NGF expression at the protein level in response to insulin exposure was also confirmed (Figures 4B and 4C). Based on these results, we focused on the roles of insulin in activating NGF expression and further investigated insulin signaling in epidermal keratinocytes. The forkhead box O1 (FOXO1) transcriptional factor, which is a key downstream target in the insulin signaling pathway in metabolic tissues, is translocated from nucleus to cytoplasm upon activation of the insulin signaling pathway (Figure 4A) ^24^. Based on this, we hypothesized that FOXO1 was responsible for insulin-dependent NGF expression in epidermal keratinocytes. Insulin-induced upregulation of NGF was associated with the translocation of FOXO1 from the nucleus to the cytoplasm (Figures 4D-4F). Moreover, *FOXO1* knockdown caused elevated NGF expression even in the absence of insulin (Figures 4G, 4H, and S3B), indicating that insulin leads to FOXO1 cytoplasmic translocation and associated de-suppression of NGF expression in epidermal keratinocytes. We then performed immunostaining of ear skin with antibodies to FOXO1 and K14, along with the pan-nuclear marker TOPRO3. Mirroring our *in vitro* assay, we found that FOXO1 was predominantly located in the nuclei of epidermal keratinocytes in controls and that there was clear cytoplasmic translocation of FOXO1 in keratinocytes of DIO skin, indicating enhanced insulin signaling in these cells (Figures 5A and 5B). Importantly, treatment with anti-PLVAP antibody reduced FOXO1 translocation, suggesting that insulin signaling is suppressed in DIO mice receiving the anti-PLVAP antibody (Figures 5A and 5B). Likewise, we found that enhanced NGF expression in keratinocytes was significantly suppressed in DIO mice receiving the anti-PLVAP antibody (Figures 5C and 5D). Notably, DIO mice with the anti-PLVAP antibody maintained increased circulating insulin, similar to DIO mice with saline, despite blocking of PLVAP (Figure S4A), suggesting that local (rather than systemic) changes in insulin delivery promotes enhanced NGF expression in the epidermal keratinocytes. Together, our findings suggest a mechanistic link between vascular permeability and pain sensation: increased vascular permeability facilitates insulin signaling activation in epidermal keratinocytes, likely by allowing insulin to diffuse from superficial dermal capillaries into the epidermis. This activation leads to insulin-mediated NGF production in keratinocytes, which in turn sensitizes sensory axons (Figure 5E).

**Figure 4.**
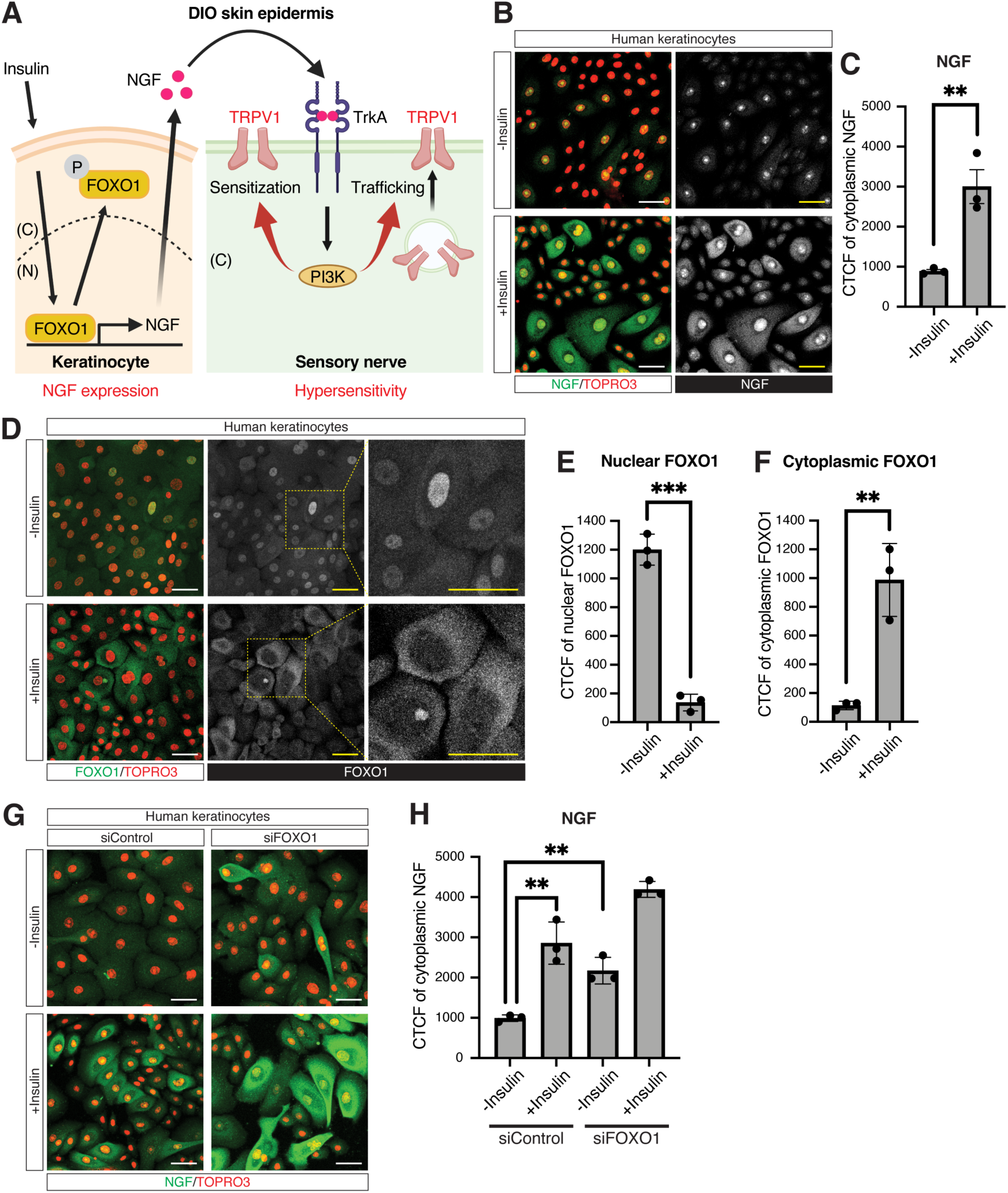
Insulin enhanced NGF expression, accompanied by FOXO1 translocation in keratinocytes. (A) Schematic diagrams illustrate the mechanisms involved in the NGF production in response to insulin signaling activation in keratinocytes (left) and the sensory hypersensitivity mediated by the NGF-TrkA-PI3K signaling pathway in sensory nerves (right). Insulin stimulates NGF expression in keratinocytes ^5^, accompanied by FOXO1 translocation from the nucleus to the cytoplasm. The NGF-TrkA-PI3K signaling pathway is involved in sensory hypersensitivity, likely by sensitizing the TRPV1 channel and facilitating its trafficking to the plasma membrane ^5^. “C” indicates the cytoplasm; “N” indicates the nucleus. (B) Representative images of immunofluorescence staining of cultured human keratinocytes, labeled with an antibody to NGF (green or white), along with the nuclear marker TOPRO3 (red), are presented. Note that non-specific nuclear signals were observed when using the anti-NGF antibody for immunostaining. Scale bars: 20 μm. (C) Quantification of cytoplasmic signals using the anti-NGF antibody staining in human keratinocytes cultured with or without insulin is shown. N=3 in each group. (D) Representative images of immunofluorescence staining of cultured human keratinocytes, labeled with an antibody to FOXO1 (green or white), along with the nuclear marker TOPRO3 (red), are presented. The dotted box regions in the middle panels are magnified in the right panels. Scale bars: 20 μm. (E) Quantification of nuclear signals using the anti-FOXO1 antibody staining in human keratinocytes cultured with or without insulin is shown. N=3 in each group. (F) Quantification of cytoplasmic signals using the anti-FOXO1 antibody staining in human keratinocytes cultured with or without insulin is shown. N=3 in each group. (G) Representative images of immunofluorescence staining of human keratinocytes cultured with or without insulin or si*FOXO1*, labeled with the anti-NGF antibody (green or white), along with TOPRO3 (red), are presented. (H) Quantification of cytoplasmic signals using the anti-NGF antibody staining in human keratinocytes cultured with or without insulin or si*FOXO1* is shown. N=3 in each group. Results are shown as the mean ± SEM. **p<0.01, ***p<0.001. P values were determined by the parametric two-tailed t test. The schematic diagrams and graphic summary were partially created with BioRender.com.

**Figure 5.**
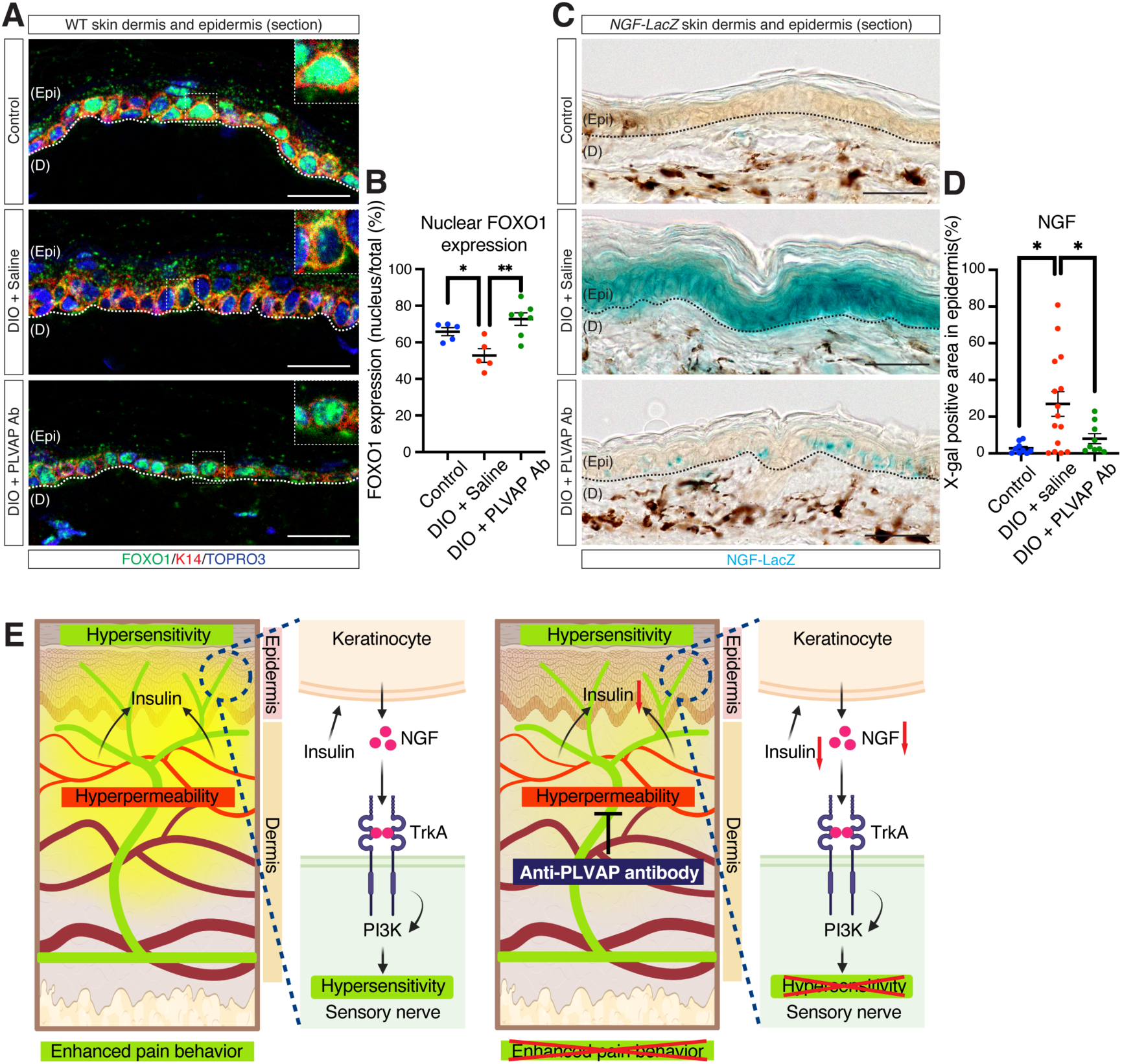
Capillary hyperpermeability leads to insulin signaling activation in epidermal keratinocytes, resulting in insulin-mediated NGF production in these cells. (A) Representative section immunohistochemical images of ear skin from control mice, DIO mice treated with saline, and DIO mice treated with the neutralizing anti-PLVAP antibody are presented. This assay uses the antibodies for FOXO1 (green), the keratinocyte marker K14 (red), along with the nuclear marker TOPRO3 (blue). Each inset displays the pattern of FOXO1 expression in a single keratinocyte. Dashed lines indicate the boundary between the epidermis and the dermis. “Epi” indicates the epidermis; “D” indicates the dermis. Scale bars: 20 μm. (B) Quantification of nuclear FOXO1 expression in keratinocytes is provided. The percentages of nuclear FOXO1 expression within the total FOXO1 expression in keratinocytes are presented. The sample sizes are as follows: N = 5 in control, N = 5 in DIO + Saline, N = 7 in DIO + PLVAP Ab. (C) Representative X-gal staining images of ear skin from *NGF-LacZ* control mice, DIO mice with saline, and DIO mice with the neutralizing anti-PLVAP antibody (blue) are presented. Dashed lines indicate the boundary between the epidermis and the dermis. Scale bars: 50 μm. (D) Quantification of the LacZ-positive area in the epidermis is provided. The sample sizes are as follows: N = 9 in control, N = 15 in DIO + Saline, N = 9 in DIO + PLVAP Ab. (E) Graphical summary illustrates how vascular hyperpermeability leads to sensory hypersensitivity in DIO skin. Increased permeability in the superficial dermal capillaries facilitates the diffusion of insulin into the epidermis, activating insulin signaling in epidermal keratinocytes. This activation leads to NGF upregulation in these keratinocytes, which in turn promotes sensory hypersensitivity in DIO skin. A neutralizing anti-PLVAP antibody reduces the diffusion of insulin, thereby decreasing NGF expression in the epidermal keratinocytes and alleviating sensory hypersensitivity. Results are shown as the mean ± SEM. *p<0.05, **p<0.01. P values were determined by the parametric two-tailed t test. The schematic diagrams and graphic summary were partially created with BioRender.com.

## Discussion

An effective therapeutic strategy for managing painful small fiber neuropathy in patients with obesity remains elusive. Due to the complexity of the condition, options to manage pain are limited to ion channel blockers, glycemic control, and lifestyle modification ^1,2,25^. In addition to obesity-associated neuropathy, treatment-induced diabetic neuropathy (TIDN), previously known as insulin neuritis, can develop as a severe, painful neuropathy following rapid glycemic control through insulin or other glucose-lowering treatments, associated with tortuous leaky blood vessels ^26–28^. However, its underlying mechanisms are not yet fully understood. We previously showed that increased NGF production in epidermal keratinocytes in response to elevated insulin levels sensitizes sensory axons, leading to hypersensitivity in obese mice where NGF directly modulates TRPV1 signaling ^5,18–23^. While blocking NGF signaling in the skin represents an interesting approach to managing peripheral pain ^5,23^, it is practically challenging to block NGF-mediated pain sensation without also affecting neuronal survival ^29,30^. In this study, we demonstrate that managing vascular hyperpermeability associated with obesity leads to a decrease in insulin-mediated NGF expression in epidermal keratinocytes, which subsequently decreases hypersensitivity. Given that PLVAP expression is confined to the endothelial fenestrae, it may represent a uniquely attractive target for managing vascular hyperpermeability and associated pain.

Skin vascular permeability is regulated and maintained by three main components: 1) paracellular junctions, such as adhesion and tight junctions; 2) transcellular pathways, which include fenestrae and caveolae; and 3) associations with pericytes ^12,31^. In the skin of DIO mice, the superficial epidermal capillaries develop endothelial fenestrae, while there are no observable changes in paracellular junctions or pericyte coverage. These endothelial fenestrae, circular openings at the EC membrane, facilitate the diffusion of small to moderately sized molecules, including peptide hormones like insulin. Recent studies have elegantly revealed the structure of PLVAP, as well as the antibody epitope in its extracellular domain ^8^. However, the exact mechanism by which the anti-PLVAP antibody blocks vascular permeability, and the potential effects of antibody treatment on endothelial fenestrae in other vascular beds remain unclear. Future experiments to establish more precise pharmacokinetics, target engagement, and mechanism of action should resolve these questions.

Endothelial fenestrations are highly dependent on VEGF-A signaling: Blocking this signaling pathway restores a non-fenestrated endothelial membrane ^32–34^, and VEGF-A regulates PLVAP in physiological and pathological conditions ^35–37^. Indeed, recent studies have effectively shown that VEGF-A plays a crucial role in maintaining fenestrated endothelium within certain EC populations in subcutaneous white adipose tissues ^38^. Further research is necessary to determine whether an excess of VEGF-A induces PLVAP expression and endothelial fenestrations in the superficial epidermal capillaries in the DIO skin.

Our studies provide the foundation for a better understanding of the link between vascular abnormalities and sensory dysfunction associated with obesity. By targeting endothelial fenestrae using PLVAP inhibitors, we can reduce sensory activity and manage neuropathic pain. Furthermore, previous studies reported that continuous activation of nociceptive sensory nerves accelerated their degeneration in obese mice ^39^, and we found that targeting endothelial fenestrae with PLVAP inhibitors helped protect against nerve degeneration. Taken together, these findings suggest that managing sensory activity by inhibiting vascular permeability during the critical window of obesity-related neuropathic pain ultimately helps prevent sensory degeneration. Thus, we propose a novel clinical approach for managing neuropathic pain associated with obesity and its potential consequent sensory decline. Additionally, it would be valuable to investigate further the potential of PLVAP inhibitors to improve vascular leakage across a wide range of disorders.

## STAR Methods

### Mice

All animal procedures were approved by the National Heart, Lung, and Blood institute (NHLBI) Animal Care and Use Committee in accordance with National Institutes of Health (NIH) research guidelines for the care and use laboratory animals. C57BL/6J mice (The Jackson Laboratory), *Pirt-GCaMP3* mice ^15^, and *NGF-LacZ* mice ^40^ were used in this study. C57BL/6J, *Pirt-GCaMP3* heterozygous, and *NGF-LacZ* heterozygous male mice were randomly separated and fed either a high-fat diet (60% fat, 20% protein, and 20% carbohydrate kcal; Research Diets, D12492) to induce DIO or a normal diet (10% fat, 20% protein, and 70% carbohydrate kcal) for control from 6 weeks-of-age with free access to water.

### Neutralizing anti-PLVAP antibody administration into DIO mice

A neutralizing anti-PLVAP antibody (clone MECA-32, BioXCell, BE0200) or a rat IgG2a isotype control (BioXCell, BE0089) was diluted to a concentration of 2.11 mg/ml in a 0.9% sodium chloride solution (Millipore Sigma, S8776). Osmotic pumps (ALZET, model 2002) were filled with the diluted antibodies according to the manufacturer’s protocol, and then surgically implanted subcutaneously into DIO mice at 20 weeks-of-age. The antibody solution was delivered systemically at a rate of 0.5 μl/hour (1.055 μg/hour) for a duration of 14 days in DIO mice.

### Pain behavior assay

The pain behavior assay was performed as described previously^5^. Briefly, the 10 μl capsaicin solution (Sigma, M2028, 0.1 mM in ethanol) was applied into the skin behind the ear after 10 minutes of habituation to stimulate sensory nerves. Wiping responses with a forelimb at the stimulus site were counted for 10 minutes.

### Ex vivo live Ca^2+^ imaging of mouse ear skin

Live Ca^2+^ imaging of mouse ear skin was performed as described previously ^5^. Ear skin was dissected from *Pirt-GCaMP3* mice, and epidermal axon fibers were imaged for 10 minutes following stimulation with 2 μM capsaicin to measure GCaMP3 fluorescent levels. The dissected ear skins were placed in a test chamber filled with synthetic interstitial fluid (SIF) ^41^ and maintained at room temperature during the Ca^2+^ imaging. To image GCaMP3 fluorescence in epidermal axon fibers within 20 μm from the skin surface, five Z-stack images were acquired every 10 seconds. Time-course images were imported into ImageJ (NIH), and Ca^2+^ transients (ΔF/F_0_) were calculated based on GCaMP3 intensities in three selected axon fibers. For statistical comparison of sensory activities, the area under the Ca^2+^ transient curve (AUC) was determined.

### Whole-mount immunostaining of mouse ear skin

Outer ear skin was dissected from mice at 22 weeks-of-age and fixed with a 4% paraformaldehyde/PBS solution at 4°C for 1 hour. Whole-mount immunostaining was performed essentially as described previously ^5,10,42^. Prior to blocking, connective tissues and fat were removed from the fixed skin. The skin tissues were incubated overnight at 4°C in a blocking buffer (0.2% TritonX-100, 10% heat inactivated goat or donkey serum in PBS), mixed with diluted primary antibodies. Staining was performed using Armenian hamster anti-PECAM-1 antibody (Millipore Sigma, MAB1398Z, 1:300), rat anti-PLVAP (clone MECA-32, BD Biosciences, 553849, 1:200), FITC-conjugated mouse anti-αSMA antibody (Sigma, F-3777, 1:250), and rat anti-PDGFRΔ antibody (Invitrogen, 16-1402-82, 1:100). For immunofluorescent detection, the skin tissues were incubated in the blocking buffer containing either Alexa-488-, Alexa-568-, or Alexa-647-conjugated secondary antibodies (Jackson ImmunoResearch or Thermo Fisher Scientific, 1:250). All confocal microscopy was carried out on a Leica TCS SP5 confocal (Leica).

### Measurement of intraepidermal nerve fiber (IENF) density

The fixed outer ear skin was immersed in a 30% sucrose/PBS solution at 4°C and then embedded in OCT compound. The tissues were cryosectioned to a thickness of 20 μm and collected on pre-cleaned slides (Fisher Scientific, 15-188-48). The skin sections were incubated overnight at 4°C in the blocking buffer with diluted primary antibodies. Staining was performed using mouse anti-Tuj1 antibody (BioLegend, 801202, 1:200) and guinea pig anti-K14 antibody (PROGEN, GP-CK14, 1:100). For immunofluorescent detection, the skin sections were incubated in the blocking buffer containing Alexa-488-and Alexa-568-conjugated secondary antibodies (Jackson ImmunoResearch or Thermo Fisher Scientific, 1:250). All confocal microscopy was carried out on a Leica TCS SP5 confocal (Leica). The IENF density was quantified by counting the number of sensory axons that crossed the epidermal-dermal junction. The linear IENF density was calculated and expressed as the number of fibers per millimeter of epidermal length (IENF/mm) for comparison.

### FOXO1 detection in epidermal keratinocytes

The fixed outer ear skin was dehydrated, embedded into paraffin wax blocks, and sectioned to a thickness of 8 μm. After deparaffinization, the skin sections were heated in a retrieval solution (DAKO, S1699) using a microwave for antigen retrieval. The skin sections were incubated overnight at 4°C in the blocking buffer with diluted primary antibodies. Staining was performed using rabbit anti-FOXO1 antibody (Cell Signaling, 2880) and guinea pig anti-K14 antibody (PROGEN, GP-CK14, 1:100). To detect FOXO1 immunofluorescence, the skin sections were incubated in the blocking buffer containing anti-rabbit HRP (PerkinElmer, NEF812001EA, 1:250), followed by incubation in Tyramide Amplification Buffer Plus (Biotium, 99832) containing OPAL 520 (Akoya Biosciences, FP1487001KT). For K14 detection, the skin sections were incubated in the blocking buffer containing Alexa-568-conjugated secondary antibody (Thermo Fisher Scientific, 1:250). Nuclei were counterstained with TOPRO3 (Invitrogen, T3605). All confocal microscopy was carried out on a Leica TCS SP5 confocal (Leica). The intensities of FOXO1 signals in K14-positive cytoplasm and TOPRO3-positive nuclei were measured with ImageJ, and then nuclear FOXO1 expression was calculated for comparison.

### X-gal staining of mouse ear skin

The outer ear skin was fixed with a 0.25% glutaraldehyde/PBS solution for 20 minutes on ice. The skin tissues were incubated overnight at 37°C in a 1 mg/ml X-gal solution. The skin tissues were washed with PBS and post-fixed with a 4% paraformaldehyde/PBS solution. For sectioning, the stained skin tissues were immersed in a 30% sucrose/PBS solution at 4°C and then embedded in OCT compound. The skin tissues were cryosectioned to a thickness of 20 μm.

### Transmission Electron Microscopy

The outer ear skin was fixed in a mixture of 2.5% glutaraldehyde and 4% paraformaldehyde in 0.1M Sodium cacodylate buffer, pH 7.4, for 60 minutes at room temperature and overnight at 4°C. The following day, the skin tissue was washed three times for ten minutes each in 0.1M Sodium cacodylate buffer, pH 7.4, and post-fixed in 1% OsO4 in 0.1M cacodylate buffer for 60 minutes on ice. Next, samples were rinsed and washed three times for 10 minutes each in water and incubated with 1% uranyl acetate overnight at 4°C. The following day samples were rinsed and washed in water and gradually dehydrated through a graded ethanol series followed by propylene oxide. Samples were then infiltrated in a mix of propylene oxide and resin (Embed 812 resin) before being infiltrated with three changes of pure resin and embedded in 100% resin and baked at 60°C for 48 hours. Ultrathin sections (65-70 nm) were cut using an ultramicrotome (Leica EM UT7), and digital micrographs were acquired on JEOL JEM 1200 EXII operating at 80Kv and equipped with AMT XR-60 digital camera. The numbers of non-fenestrated endothelial cells and fenestrated endothelial cells were manually counted for comparison. The distance between endothelial cells were measured using ImageJ to assess junction stability.

### Vascular permeability assay by intravital live imaging of ear skin

Mice were anesthetized using isoflurane and placed on a heating-plate on the stage of Leica SP8 two-photon microscope (Leica). The ear was positioned on a customized imaging stage and immersed in imaging gel. DyLight 649-conjugeted Lycopersicon Esculentum (Tomato) lectin was injected through the cannulated tail vein to visualize blood vessels. We focused on the capillary vessels in the superficial dermis based on the lectin signals. Additionally, tetramethylrhodamine-conjugated 40 kDa dextran was injected through the tail vein, and capillary vessels were imaged every minute to measure dextran leakage. Time-course Z-stack images were imported into Fiji, and Fast4Dreg plugin was used for minor drift correction. The capillary vessels were manually selected to exclude large-diameter blood vessels and further segmented using Labkit plugin in Fiji. In the dextran 40 kDa channel, dextran signals inside the capillary vessels were removed, and then dextran signals outside the capillary vessels were quantified every minute. The dextran intensity outside the capillary vessels was analyzed for comparison. To confirm dextran leakage, fluorescence lifetime imaging microscopy was performed before and after the dextran injection.

### Human keratinocyte culture

To prevent insulin signaling activation, insulin was omitted from the medium. Human keratinocytes (ScienCell, 2100) were cultured in the KGM Glod Keratinocyte Growth Medium BulletKit (Lonza, 00192060), with or without 400 nM insulin (Thermo Fisher Scientific, 12585014), for 3-4 days. 3 days prior to insulin stimulation, either scramble control (Sigme, SIC001) or *FOXO1* siRNA (EHU156591) was transfected into human keratinocytes using Lipofectamine 3000 Transfection Reagent (Thermo Fisher Scientific, L3000008). Cells were fixed with a 4% paraformaldehyde/PBS solution for 30 minutes and incubated overnight at 4°C in the blocking buffer with diluted primary antibodies.

Staining was performed using rabbit anti-NGF antibody (abcam, ab52918, 1:300) and rabbit anti-FOXO1 antibody (Cell Signaling, 2880, 1:100). For immunofluorescent detection, cells were incubated in the blocking buffer containing Alexa-488-conjugated secondary antibody (Jackson ImmunoResearch, 1:250). Nuclei were counterstained with TOPRO3 (Invitrogen, T3605). All confocal microscopy was carried out on a Leica TCS SP5 confocal (Leica). The expression of NGF and FOXO1 was calculated with corrected total cell fluorescence (CTCF) as the mean fluorescence value after subtracting the background signals.

### Quantitative RT-PCR

Total RNAs from cultured human keratinocytes were extracted using RNeasy Mini kits (QUAGEN) according to the manufacturer’s protocol. Residual genomic DNA was digested with Ambion DNase I (Thermo Fisher Scientific, AM2222). First-strand cDNA was synthesized using the SuperScript III First-Strand Synthesis SuperMix (Thermo Fisher Scientific, 18080400). Quantitative RT-PCR was performed using THUNDERBIRD Next SYBR qPCR Mix (TOYOBO). All data of quantitative RT-PCR were calculated using the ddCt method with *GAPDH* as normalization controls. Primers are listed in Table S1.

### Statistical analysis

Results were presented as the mean ± SEM. P values were determined by the parametric two-tailed t test. p < 0.05 was considered statistically significant.

## Resource availability

### Lead contact

Requests for further information and resources should be directed to and will be fulfilled by the lead contact, Yoh-suke Mukouyama.

### Materials availability

This study did not generate new unique reagents.

### Data and code availability

This study does not generate datasets.

This study does not report original code.

Any additional information required to reanalyze the data reported in this paper is available from the lead contact upon request.

## Acknowledgements

Thanks to T. Clark and the staff of the NIH Bldg50 Animal Facility for assistance with mouse breeding and care; Z. A. Syed of the NHLBI Electron Microscopy Core for transmission electron microscopy; H. Alkaissi of the NDDK, T. Arnold of the UCSF, K. Campbell of CCHMC, and V. Bautch of UNC for valuable discussion and manuscript editing; K. Gill for laboratory management, technical support and manuscript editing; J. Dawes and S. Thacker for administrative assistance. Thanks also to members of Laboratory of Stem Cell and Neuro-Vascular Biology for thoughtful discussion and technical suggestions. This work was supported by the Intramural Research Program of the National Heart, Lung, and Blood Institute, National Institutes of Health (HL005702-20 to Y.M.). The contributions of the NIH authors were made as part of their official duties as NIH federal employees, are in compliance with agency policy requirements, and are considered Works of the United States Government. However, the findings and conclusions presented in this paper are those of the authors and do not necessarily reflect the views of the NIH or the U.S. Department of Health and Human Services.

## Author contributions

Conceptualization: Y.K., Y-S.M.; Formal analysis: Y.K., J.R.S, S.S., C.A.C; Funding acquisition: Y-S.M.; Investigation: Y.K., J.R.S, S.S., C.A.C; Methodology: Y.K., J.R.S, S.S., C.A.C; Project administration: Y-S.M.; Resources: C.A.C; Supervision: Y-S.M.; Validation: Y.K., J.R.S, S.S., C.A.C; Visualization: Y.K., J.R.S, S.S., C.A.C; Writing - original draft: Y.K., Y-S.M.; Writing - review & editing: Y.K., J.R.S, S.S., C.A.C, Y-S.M.

## References

1. Feldman, E.L., Callaghan, B.C., Pop-Busui, R., Zochodne, D.W., Wright, D.E., Bennett, D.L., Bril, V., Russell, J.W., and Viswanathan, V. (2019). Diabetic neuropathy. Nature Reviews Disease Primers 5, 41. 10.1038/s41572-019-0092-1.

2. Vincent, A.M., Callaghan, B.C., Smith, A.L., and Feldman, E.L. (2011). Diabetic neuropathy: cellular mechanisms as therapeutic targets. Nature Reviews Neurology 7, 573–583. 10.1038/nrneurol.2011.137.

3. Graupera, M., and Claret, M. (2018). Endothelial Cells: New Players in Obesity and Related Metabolic Disorders. Trends in Endocrinology & Metabolism 29, 781–794. 10.1016/j.tem.2018.09.003.

4. Stapleton, P.A., James, M.E., Goodwill, A.G., and Frisbee, J.C. (2008). Obesity and vascular dysfunction. Pathophysiology 15, 79–89. 10.1016/j.pathophys.2008.04.007.

5. Koui, Y., Song, S., Dong, X., and Mukouyama, Y.-s. (2025). Local keratinocyte-nociceptor interactions enhance obesity-mediated painful small fiber neuropathy via NGF-TrkA-PI3K signaling axis. iScience 28. 10.1016/j.isci.2025.112047.

6. Stan, R.V., Kubitza, M., and Palade, G.E. (1999). PV-1 is a component of the fenestral and stomatal diaphragms in fenestrated endothelia. Proc Natl Acad Sci U S A 96, 13203–13207. 10.1073/pnas.96.23.13203.

7. Mou, X., Leeman, S.M., Roye, Y., Miller, C., and Musah, S. (2024). Fenestrated Endothelial Cells across Organs: Insights into Kidney Function and Disease. International Journal of Molecular Sciences 25, 9107.

8. Chang, T.-H., Hsieh, F.-L., Gu, X., Smallwood, P.M., Kavran, J.M., Gabelli, S.B., and Nathans, J. (2023). Structural insights into plasmalemma vesicle-associated protein (PLVAP): Implications for vascular endothelial diaphragms and fenestrae. Proceedings of the National Academy of Sciences 120, e2221103120. doi:10.1073/pnas.2221103120.

9. Denzer, L., Muranyi, W., Schroten, H., and Schwerk, C. (2023). The role of PLVAP in endothelial cells. Cell Tissue Res 392, 393–412. 10.1007/s00441-023-03741-1.

10. Yamazaki, T., Li, W., Yang, L., Li, P., Cao, H., Motegi, S.I., Udey, M.C., Bernhard, E., Nakamura, T., and Mukouyama, Y.S. (2018). Whole-Mount Adult Ear Skin Imaging Reveals Defective Neuro-Vascular Branching Morphogenesis in Obese and Type 2 Diabetic Mouse Models. Sci Rep 8, 430. 10.1038/s41598-017-18581-7.

11. Egawa, G., Nakamizo, S., Natsuaki, Y., Doi, H., Miyachi, Y., and Kabashima, K. (2013). Intravital analysis of vascular permeability in mice using two-photon microscopy. Scientific Reports 3, 1932. 10.1038/srep01932.

12. Ono, S., Egawa, G., and Kabashima, K. (2017). Regulation of blood vascular permeability in the skin. Inflammation and Regeneration 37, 11. 10.1186/s41232-017-0042-9.

13. Claesson-Welsh, L., Dejana, E., and McDonald, D.M. (2021). Permeability of the Endothelial Barrier: Identifying and Reconciling Controversies. Trends in Molecular Medicine 27, 314–331. 10.1016/j.molmed.2020.11.006.

14. Nakagami, Y., Hatano, E., Chayama, Y., and Inoue, T. (2019). An anti-PLVAP antibody suppresses laser-induced choroidal neovascularization in monkeys. European Journal of Pharmacology 854, 240–246. 10.1016/j.ejphar.2019.04.035.

15. Kim, Y.S., Chu, Y., Han, L., Li, M., Li, Z., LaVinka, P.C., Sun, S., Tang, Z., Park, K., Caterina, M.J., et al. (2014). Central terminal sensitization of TRPV1 by descending serotonergic facilitation modulates chronic pain. Neuron 81, 873–887. 10.1016/j.neuron.2013.12.011.

16. Pan, H., Fatima, M., Li, A., Lee, H., Cai, W., Horwitz, L., Hor, C.C., Zaher, N., Cin, M., Slade, H., et al. (2019). Identification of a Spinal Circuit for Mechanical and Persistent Spontaneous Itch. Neuron 103, 1135–1149.e1136. 10.1016/j.neuron.2019.06.016.

17. Shimada, S.G., and LaMotte, R.H. (2008). Behavioral differentiation between itch and pain in mouse. Pain 139, 681–687. 10.1016/j.pain.2008.08.002.

18. Barker, P.A., Mantyh, P., Arendt-Nielsen, L., Viktrup, L., and Tive, L. (2020). Nerve Growth Factor Signaling and Its Contribution to Pain. J Pain Res 13, 1223–1241. 10.2147/jpr.S247472.

19. Chang, D.S., Hsu, E., Hottinger, D.G., and Cohen, S.P. (2016). Anti-nerve growth factor in pain management: current evidence. J Pain Res 9, 373–383. 10.2147/jpr.S89061.

20. Denk, F., Bennett, D.L., and McMahon, S.B. (2017). Nerve Growth Factor and Pain Mechanisms. Annu Rev Neurosci 40, 307–325. 10.1146/annurev-neuro-072116-031121.

21. Lawrence, G.W., Zurawski, T.H., and Dolly, J.O. (2021). Ca(2+) Signalling Induced by NGF Identifies a Subset of Capsaicin-Excitable Neurons Displaying Enhanced Chemo-Nociception in Dorsal Root Ganglion Explants from Adult pirt-GCaMP3 Mouse. Int J Mol Sci 22. 10.3390/ijms22052589.

22. Stratiievska, A., Nelson, S., Senning, E.N., Lautz, J.D., Smith, S.E., and Gordon, S.E. (2018). Reciprocal regulation among TRPV1 channels and phosphoinositide 3-kinase in response to nerve growth factor. Elife 7. 10.7554/eLife.38869.

23. Cheng, H.T., Dauch, J.R., Hayes, J.M., Hong, Y., and Feldman, E.L. (2009). Nerve Growth Factor Mediates Mechanical Allodynia in a Mouse Model of Type 2 Diabetes. Journal of Neuropathology & Experimental Neurology 68, 1229–1243. 10.1097/NEN.0b013e3181bef710.

24. Teaney, N.A., and Cyr, N.E. (2023). FoxO1 as a tissue-specific therapeutic target for type 2 diabetes. Front Endocrinol (Lausanne) 14, 1286838. 10.3389/fendo.2023.1286838.

25. Jensen, T.S., Backonja, M.-M., Jiménez, S.H., Tesfaye, S., Valensi, P., and Ziegler, D. (2006). New perspectives on the management of diabetic peripheral neuropathic pain. Diabetes & Vascular Disease Research 3, 108–119. 10.3132/dvdr.2006.013.

26. Gibbons, C.H. (2017). Treatment-Induced Neuropathy of Diabetes. Curr Diab Rep 17, 127. 10.1007/s11892-017-0960-6.

27. Gibbons, C.H. (2020). Treatment induced neuropathy of diabetes. Auton Neurosci 226, 102668. 10.1016/j.autneu.2020.102668.

28. Tesfaye, S., Malik, R., Harris, N., Jakubowski, J.J., Mody, C., Rennie, I.G., and Ward, J.D. (1996). Arterio-venous shunting and proliferating new vessels in acute painful neuropathy of rapid glycaemic control (insulin neuritis). Diabetologia 39, 329–335. 10.1007/bf00418349.

29. Apfel, S.C., Schwartz, S., Adornato, B.T., Freeman, R., Biton, V., Rendell, M., Vinik, A., Giuliani, M., Stevens, J.C., Barbano, R., and Dyck, P.J. (2000). Efficacy and safety of recombinant human nerve growth factor in patients with diabetic polyneuropathy: A randomized controlled trial. rhNGF Clinical Investigator Group. Jama 284, 2215–2221. 10.1001/jama.284.17.2215.

30. Bramson, C., Herrmann, D.N., Carey, W., Keller, D., Brown, M.T., West, C.R., Verburg, K.M., and Dyck, P.J. (2015). Exploring the Role of Tanezumab as a Novel Treatment for the Relief of Neuropathic Pain. Pain Medicine 16, 1163–1176. 10.1111/pme.12677.

31. Goddard, L.M., and Iruela-Arispe, M.L. (2013). Cellular and molecular regulation of vascular permeability. Thromb Haemost 109, 407–415. 10.1160/th12-09-0678.

32. Yang, Y., Zhang, Y., Cao, Z., Ji, H., Yang, X., Iwamoto, H., Wahlberg, E., Länne, T., Sun, B., and Cao, Y. (2013). Anti-VEGF- and anti-VEGF receptor-induced vascular alteration in mouse healthy tissues. Proc Natl Acad Sci U S A 110, 12018–12023. 10.1073/pnas.1301331110.

33. Roberts, W.G., and Palade, G.E. (1995). Increased microvascular permeability and endothelial fenestration induced by vascular endothelial growth factor. J Cell Sci 108 ( *Pt 6*), 2369–2379. 10.1242/jcs.108.6.2369.

34. Kamba, T., Tam, B.Y., Hashizume, H., Haskell, A., Sennino, B., Mancuso, M.R., Norberg, S.M., O’Brien, S.M., Davis, R.B., Gowen, L.C., et al. (2006). VEGF-dependent plasticity of fenestrated capillaries in the normal adult microvasculature. Am J Physiol Heart Circ Physiol 290, H560–576. 10.1152/ajpheart.00133.2005.

35. Fan, Z., Karakone, M., Nagarajan, S., Nagy, N., Mildenberger, W., Petrova, E., Hinte, L.C., Bijnen, M., Häne, P., Nelius, E., et al. (2024). Macrophages preserve endothelial cell specialization in the adrenal gland to modulate aldosterone secretion and blood pressure. Cell Rep 43, 114395. 10.1016/j.celrep.2024.114395.

36. Kim, S.A., Kim, S.J., Choi, Y.A., Yoon, H.J., Kim, A., and Lee, J. (2020). Retinal VEGFA maintains the ultrastructure and function of choriocapillaris by preserving the endothelial PLVAP. Biochem Biophys Res Commun 522, 240–246. 10.1016/j.bbrc.2019.11.085.

37. Strickland, L.A., Jubb, A.M., Hongo, J.A., Zhong, F., Burwick, J., Fu, L., Frantz, G.D., and Koeppen, H. (2005). Plasmalemmal vesicle-associated protein (PLVAP) is expressed by tumour endothelium and is upregulated by vascular endothelial growth factor-A (VEGF). J Pathol 206, 466–475. 10.1002/path.1805.

38. Hasan, S.S., John, D., Rudnicki, M., AlZaim, I., Eberhard, D., Moll, I., Taylor, J., Klein, C., von Heesen, M., Conradi, L.-C., et al. (2025). Obesity drives depot-specific vascular remodeling in male white adipose tissue. Nature Communications 16, 5392. 10.1038/s41467-025-60910-2.

39. Jayaraj, N.D., Bhattacharyya, B.J., Belmadani, A.A., Ren, D., Rathwell, C.A., Hackelberg, S., Hopkins, B.E., Gupta, H.R., Miller, R.J., and Menichella, D.M. (2018). Reducing CXCR4-mediated nociceptor hyperexcitability reverses painful diabetic neuropathy. J Clin Invest 128, 2205–2225. 10.1172/jci92117.

40. Liu, Y., Rutlin, M., Huang, S., Barrick, C.A., Wang, F., Jones, K.R., Tessarollo, L., and Ginty, D.D. (2012). Sexually dimorphic BDNF signaling directs sensory innervation of the mammary gland. Science 338, 1357–1360. 10.1126/science.1228258.

41. Zimmermann, K., Hein, A., Hager, U., Kaczmarek, J.S., Turnquist, B.P., Clapham, D.E., and Reeh, P.W. (2009). Phenotyping sensory nerve endings in vitro in the mouse. Nat Protoc 4, 174–196. 10.1038/nprot.2008.223.

42. Yamazaki, T., Li, W., and Mukouyama, Y.S. (2018). Whole-mount Confocal Microscopy for Adult Ear Skin: A Model System to Study Neuro-vascular Branching Morphogenesis and Immune Cell Distribution. J Vis Exp. 10.3791/57406.

