## Supplementary material for "OBESITY-INDUCED ENDOTHELIAL FENESTRATION AND CAPILLARY LEAKAGE CONTRIBUTE TO INCREASED PAIN SENSATION": Figure S1-S4 and Table S1

**Figure S1, related to Figure. 1:** Adhesion junctions and pericyte coverage are maintained in the skin capillaries of DIO mice.

**Figure S2, related to Figure 3:** PLVAP inhibitor manages pain sensation in DIO skin.

**Figure S3, related to Figure 4:** Insulin leads to NGF expression in human keratinocytes in culture.

**Figure. S4, related to Figure 5:** DIO mice with the anti-PLVAP antibody maintained increased circulating insulin.

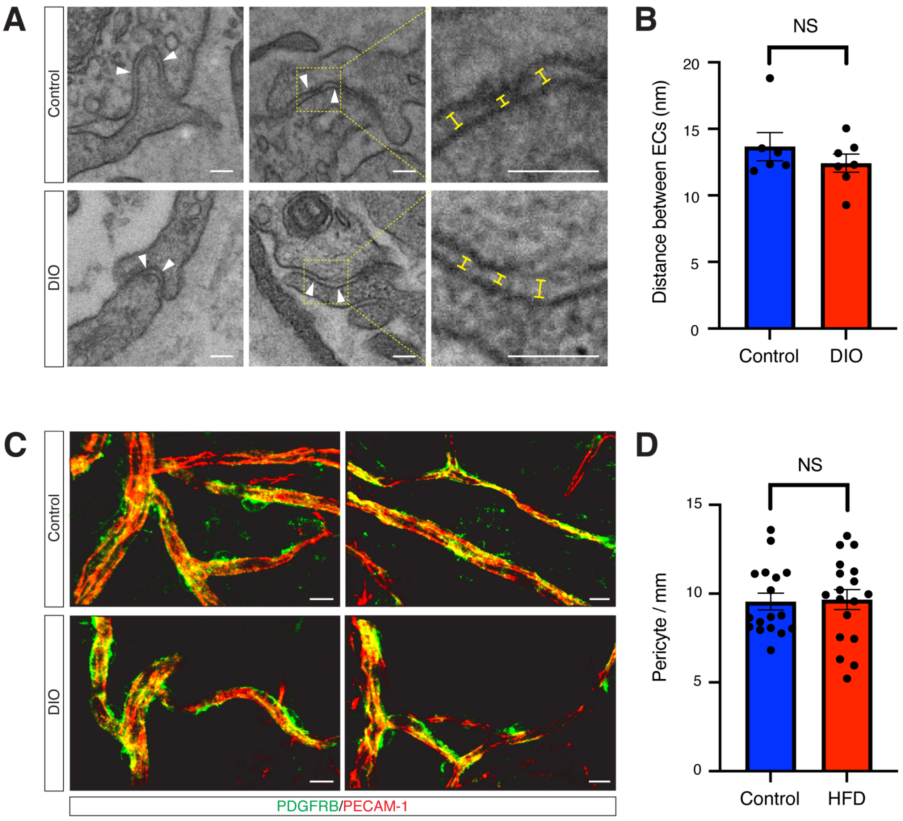


**Figure S1. Adhesion junctions and pericyte coverage are maintained in the skin capillaries of DIO mice.**

(A) Representative transmission electron microscopy images depict capillary vessels in the superficial dermis of both control and DIO mice. Arrowheads indicate endothelial junctions and the dotted box regions in the middle panels are magnified in the right panels. The distances between endothelial cells (ECs) were measured to assess the stability of EC-adhesion junctions, as indicated by the yellow lines. Scale bars: 100 nm. (B) Quantification of the distances between capillary endothelial cells is shown. The sample sizes are as follows: N = 6 in control, N = 7 in DIO. (C) Representative whole-mount double immunofluorescence images of the superficial dermal vasculature in the ear skin of control and DIO mice at 22 weeks-of-age, labeled with antibodies to the pericyte marker PDGFRβ (green) and the pan-endothelial cell marker PECAM-1 (red) are presented. Scale bars: 10 μm. (D) Quantification of pericyte coverage is shown. The sample sizes are as follows: N = 17 in control, N = 18 in DIO.

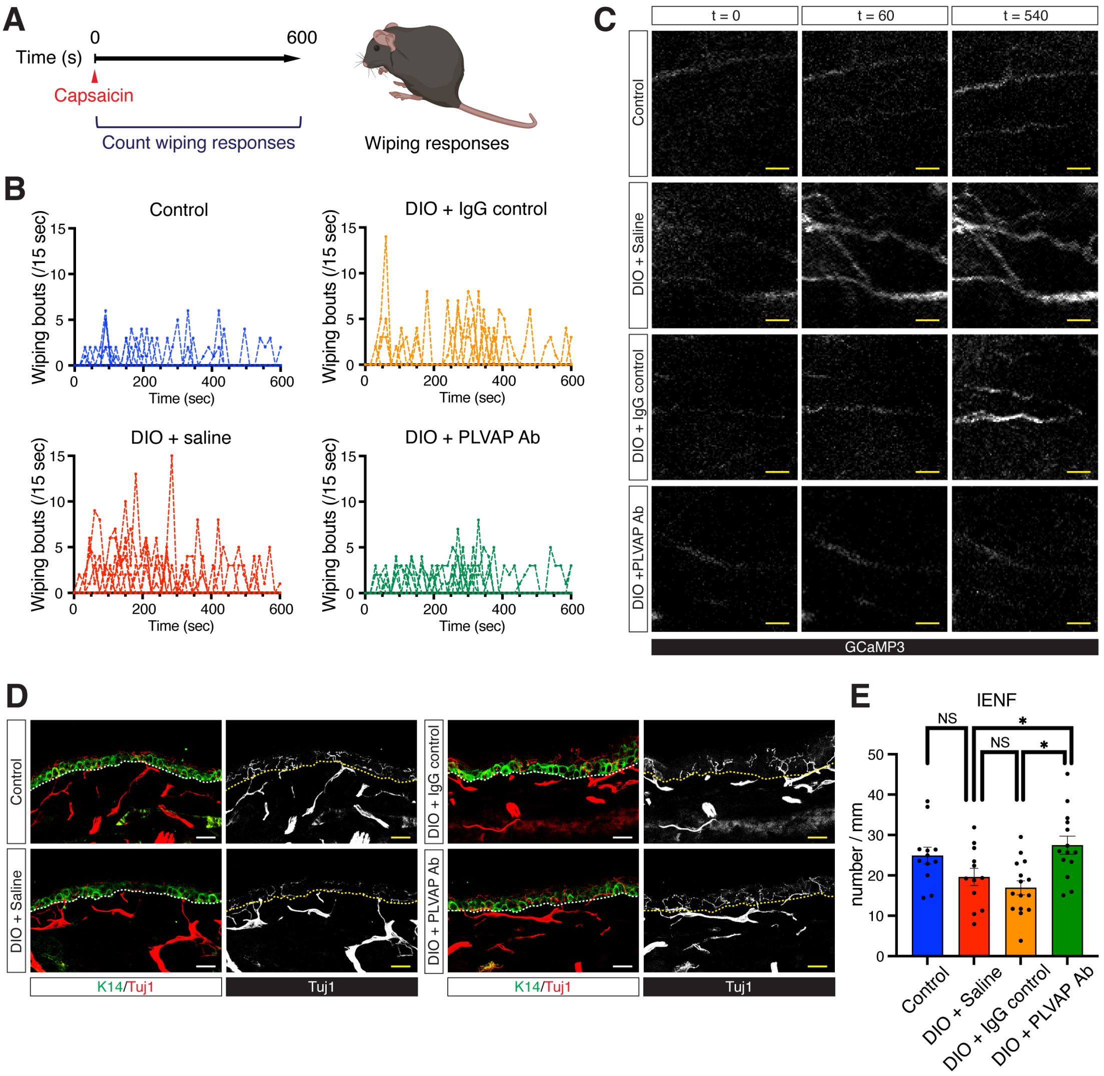


**Figure S2. PLVAP inhibitor manages pain sensation in DIO skin.**

(A) Schematic diagram illustrates the capsaicin-mediated acute pain behavior assay conducted on mouse ear skin. Forelimb wiping responses were recorded for 600 seconds (10 minutes) after capsaicin was applied to the skin behind the ears. (B) Forelimb wiping responses following capsaicin application are shown for *Pirt-GCaMP3* mice on a control diet (control, blue), *Pirt-GCaMP3* mice with DIO receiving saline (DIO + Saline, red), *Pirt-GCaMP3* mice with DIO receiving IgG control (DIO + IgG control, yellow), and *Pirt-GCaMP3* mice with DIO receiving the anti-PLVAP antibody (DIO + PLVAP Ab, green). Each dot represents a wiping bout recorded every 15 seconds. (C) Representative GCaMP3 images (white) in a single axon within ear skin explants from *Pirt-GCaMP3* mice (control), DIO + Saline, DIO + IgG control, and DIO + PLVAP Ab. GCaMP3 images are presented at t = 0, 60, and 540 seconds. Scale bars: 20 μm. (D) Representative images of section double immunofluorescence staining of the ear skin of *Pirt-GCaMP3* mice (control), DIO + Saline, DIO + IgG control, and DIO + PLVAP Ab, labeled with antibodies to the keratinocyte marker K14 (green) and the pan-axon/neuron marker neuron-specific class III b-tubulin (Tuj1, red) are presented. Dashed lines indicate the border between the epidermis and the dermis. Scale bars: 20 μm. (E) Quantification of intraepidermal nerve fibers (IENF) in each image is shown. The linear IENF density is calculated and expressed as the number of fibers per millimeter of epidermal length (IENF/mm). Results are shown as the mean ± SEM. $*p<0.05$; NS, not significant ($p > 0.05$). *P* values were determined by the parametric two-tailed *t* test.

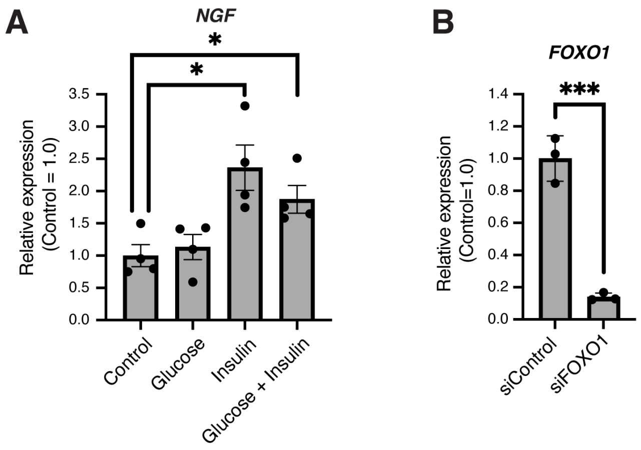


**Figure S3. Insulin leads to NGF expression in human keratinocytes in culture.**
(A) Relative *NGF* mRNA expression levels in human keratinocytes cultured with glucose, insulin, or both are shown. These expression levels are normalized to those in human keratinocytes cultured without glucose and insulin. N=4 in each group. (B) *FOXO1* knockdown efficiency was evaluated by measuring relative *FOXO1* mRNA expression levels in human keratinocytes transfected with either a scramble control siRNA (siControl) or a *FOXO1* siRNA (si*FOXO1*). These expression levels are normalized to those in human keratinocytes transfected with siControl. N=3 in each group. Results are shown as the mean ± SEM. *p<0.05, ***p<0.001. P values were determined by the parametric two-tailed t test.

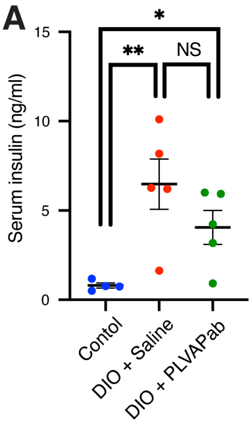


**Figure S4. DIO mice with the anti-PLVAP antibody maintained increased circulating insulin.**

(A) Serum insulin levels for *Pirt-GCaMP3* mice (control), DIO + Saline, and DIO + PLVAP Ab are shown. Results are shown as the mean ± SEM. *p<0.05, *p<0.01; NS, not significant (p > 0.05). P values were determined by the parametric two-tailed t test.

**Table S1. List of quantitative RT-PCR primers for mouse genes.**

| | Forward sequence | Reverse sequence |
|---|---|---|
| *GAPDH* | GTCTCCTCTGACTTCAACAGCG | ACCACCCTGTTGCTGTAGCCAA |
| *NGF* | ACCCGCAACATTACTGTGGACC | GACCTCGAAGTCCAGATCCTGA |
| *FOXO1* | CTACGAGTGGATGGTCAAGAGC | CCAGTTCCTTCATTCTGCACACG |
